# Proteolytic Analysis of Epsilon 34 Phage Tailspike protein indicates Partial Sensitivity to Proteinase K

**DOI:** 10.1101/2022.04.12.488085

**Authors:** Joseph A. Ayariga, Robert Villafane

## Abstract

Purified bacteriophage ε34 tailspike protein (ε34 TSP) can bind to *Salmonella newington* (S. *newington*) via the binding site of the protein, which is the O antigens of the LPS of the bacterium. We demonstrated elsewhere that purified ε34 TSP possessed bacteria lytic property on S. *newington*. The ε34 TSP has been shown via computational prediction to consist of parallel β-helices like that of P22 TSP. These protein moieties are among the simplest repetitive structural elements in proteins. There exist extensive research on the folding behavior of β-helix proteins, which also provides insight on how amyloid fibrils are generated since these proteins consist of similar parallel β-helix motifs. One of the most significantly studied system for investigating protein folding is the from the *Salmonella* bacteriophage P22. The major component of this protein is a right-handed parallel β-helix with 13 rungs. Initial *in silico* analysis of the ε34 phage TSP indicates similar structural similarity to the P22 TSP. Our previous studies indicated that despite the similarities of the two proteins, P22 TSP shows higher resistance to proteases (e.g. trypsin) and heat compared to ε34 TSP. In this study we further proof that ε34 TSP is partially sensitive to proteinase K, whereas P22 TSP is completely resistant to this protein. Detailed analysis indicates that specific structural motifs of ε34 TSP is insensitive to the protease, whereas other regions of the protein showed susceptibility to it.

## 1. Introduction

*Salmonella* has been demonstrated to cause several acute and chronic infections in humans **[1]**. These bacteria is known to be responsible for over 1.2 million illnesses in the US each year **[2, 3, 4, 5]**. Salmonella species are prone to acquiring resistance to various classes of antibiotics this sparks the need to look for methods of treating *salmonella* related infections other than the use of antibiotics, thus calling for the application of bacteriophages to manage, control and treat bacterial infections **[6, 7]**. Several bacteriophages specific to *Salmonella* have been studied **[8, 9, 10]**. ε34 phage belongs to the P22-like phages **[11, 12, 13]**. The of ε34 phage has been demonstrated to possesses similar functional and structural characteristics as the P22 phage TSP **[10]**. The predicted structure of Epsilon 34 TSP consists of a globular head binding domain, a solenoid-shape parallel beta-helix domain that is used in the LPS binding process as also found in the P22 phage TSP **[9, 14, 15]**. Phages or phage derived components can complement host cell defense against bacteria **[3, 16]**.

Purified bacteriophage ε34 TSP can bind to *S. newington* via the binding site of the protein, which is the O antigens of the LPS of the bacterium **[9]**. We demonstrated elsewhere that purified ε34 TSP possessed bacteria lytic property on *S. newington* **[9, 17]**. The ε34 TSP has been shown via computational prediction to consist of parallel β- helices like that of P22 TSP **[9]**. These protein moieties are among the simplest repetitive structural elements in proteins **[18]**. There exist extensive research on the folding behavior of β-helix proteins, which also provides insight on how amyloid fibrils are generated since these proteins consist of parallel β-helix motifs **[19, 20, 21, 22]**. One of the most significantly studied system for investigating protein folding is the from the Salmonella bacteriophage P22 **[3, 9, 23]**. The major component of this protein is a right-handed parallel β-helix with 13 rungs **[24]**. Initial *in silico* analysis of the ε34 phage indicates similar structural similarity to the P22 TSP **[9]**. Our previous studies indicated that despite the similarities of the two proteins, P22 TSP shows higher resistance to proteases (e.g. trypsin) and heat compared to ε34 TSP **[3]**. Several common proteases that have been used in most published works include chymotrypsin, trypsin, enterokinase, proteinase K, factor Xa, Staphylococcus aureus protease, papain etc., in proteolytic analysis [25, 26, 27, 28, 29], and their data are demonstrated to be consistent with expected fragments in denatured substrates. In native protein, protein structural states at which proteins still retains their tertiary conformations, most enzymes sites are shielded and hence inaccessible to the enzyme thereby imparting resistance to the enzyme, however, if there exist structural domains that are flexible, or linker regions that are solvent accessible, these areas can easily be attacked by the enzymes leading to specific cleavages and production of truncated products with unique electrophoretic mobility.

Also known as protease K or endopeptidase K, or Tritirachium alkaline proteinase, or Tritirachium album proteinase is a broad-spectrum serine protease that is capable of digesting keratin, hence the “name proteinase K”. Belonging to the S8 family of peptidase, it has a molecular weight of 28.9kDa. Unlike trypsin, proteinase K goes after the peptide bond adjacent to the carboxyl group of aliphatic and aromatic amino acids with blocked alpha amino groups. This protease is stable over a wide pH range (4–12), with a pH optimum of pH 8.0 **[30]**.

Our approach here was to unravel the native Eε34 TSP while exposing it carefully and gradually to proteinase K. To allow for comparability, samples of the s were subjected to proteolysis under non-denaturing conditions first, while to another sample, the gradual denaturation via heating at 70 °C and 80 °C was employed for predetermined time points and aliquots withdrawn at these time points and treated with the enzyme. Hence, the peptide maps obtained when the proteins were completely folded and hence least perturbed by the protease, juxtaposed with peptide maps generated during denaturing conditions as well as complete unfolded protein samples could be compared. Thus, higher structure information such as thermostable domains could be deciphered and their stability against heat as well as unfolding rates of the truncated products revealed.

Thus, in this study, we further proved that ε34 TSP is partially sensitive to proteinase K, whereas P22 TSP is completely resistant to this protein. Detailed analysis indicates that specific structural motifs of ε34 TSP is insensitive to the protease, whereas other regions of the protein showed susceptibility to it.

## 2. Materials and Methods

The cloning and protein expression, as well as the initial thermal characterization of Eε34 TSP have been documented previously in our laboratory **[9]**.

### 2.1.1. *In silico* disorder region prediction and PK sites

The entire amino acid sequence of the wild-type and the Eε34 TSP were submitted to the online Predictor of Natural Disordered Regions server [http://www.pondr.com/] as well as the secondary structure prediction server known as RaptorX [http://raptorx.uchicago.edu, **31]**. Predictions depicted amino acids that are in disordered regions as red and those in ordered region as blue. Prediction of PK cleave sites were carried out using the online programs PeptideCutter from Expasy (https://www.expasy.org/resources/peptidecutter) that predicts cleavage sites in the protein’s primary sequence.

### 2.2. Proteinase K treatment

The proteinase K treatment process followed a similar protocol used by Bosch et al., 2003 **[29]**. Stock samples of Eε34, MEε34 TSP in solutions (10 mg/mL) were diluted to the appropriate concentrations of 0.5 mg/mL using tris-HCl buffer at pH 7.4. The protein samples were then incubated for specific time points and at the specified temperatures as described below and finally, were subjected to proteinase K digestion. Samples reactions were immediately stopped by the addition of protease inhibitor, then subjected to either Native PAGE or SDS-PAGE analysis.

### 2.3. Eε34 TSP sensitivity to Proteinase K under 2% SDS at physiological temperature

To investigate the structural dynamics of our protein under slightly denaturing conditions, we deployed the use of the denaturing detergent, sodium dodecyl sulfate (SDS), and denatured the protein samples for 6 h. A sample of 0.5 mg/ml of Eε34 and Mε34 TSP were dialyzed against neutral buffer consisting of 50 mg/ml tris-HCl, pH 7.4, this was followed by mixing with 2% SDS then treated with the proteinase K enzyme at 1:20 w/w of enzyme to protein sample. The treated samples were then incubated in a water bath at 37 °C for 6 h. Afterward, the proteolytic reactions were stopped by the addition of protease inhibitor at a 10 µg/ml final concentration and the samples were subjected to electrophoretic analysis using native polyacrylamide gel. Placebos received buffer instead of the enzyme and heated controls samples also received no treatment at all except heating of these samples in 2% SDS loading buffer at 100 °C for 5 min. Samples were run at 100 volts for 250 min under non-denaturing running buffer conditions. Gels were then visualized via Coomassie blue staining, and the gels photographed using ChemiDoc XRS installed with Quantity One.

### 2.4. Eε34 sensitivity to Proteinase K under increasing concentrations of enzyme

Concentration dependent tryptic proteolysis of Eε34 TSP at 37 °C via a dose dependent treatment of our TSP to trypsin was carried out. In this work, we measured the ability of the enzyme to digest Eε34 TSP in increasing concentrations of the enzyme (trypsin). 1 mg/ml of Eε34 TSP samples were initially dialyzed against buffer consisting of 50 mg/ml tris-HCl, pH 7.4 to remove protease inhibitors then mixed with trypsin in differing ratios. A final concentration of 100 µg, 200 µg, 300 µg, 400 µg and 500 µg of trypsin were used to treat 1 mg of Eε34 TSP. Thus, a protein to enzyme ratio of 100:1, 100:2, 100:3, 100:4, 100:5 respectively for each treatment sample were achieved. Then samples were incubated for 6 h under 37 °C. Afterwards samples were run in 7.5% polyacrylamide gel and the gels photographed using ChemiDoc XRS installed with Quantity One. Experiments were repeated thrice and the densitometric values of the resulting gels were recorded via Image J 1.52 software and values plotted into graphs.

### 2.5. Eε34 TSP sensitivity to Proteinase K under protease-heat combination

To learn of the unfolding kinetics the protein and the structural stabilities of the various domains that constitute the protein, we subjected samples of the protein to heat at 70 °C, or 80 °C in the absence of denaturants, and aliquots withdrawn and quenched at specified time points (0, 2.5, 5, 10, 15, 20, 30 and 60 min). A 1:20 w/w ratio of proteinase K to Eε34 TSP sample was used for the proteolytic process. The enzyme digestion was allowed for 30 min, and quenching was achieved via mixing reaction samples with a protease inhibitor followed by the addition of SDS-free gel loading buffer consisting of 50 mM Tris-HCl, 25% glycerol, and 0.01% bromophenol blue, and SDS free running buffer consisting of 25 mM Tris-HCl, and 0.2 M glycine. Samples of triplicate experiments were electrophoresed for 60 min and stained with Coomassie blue, and then intensities of the bands that were generated were then recorded. The densitometric values were compiled from three replicate experiments and averaged. The mean densitometric values for the 70 °C and 80 °C heating experiments were plotted against time of heating.

## Statistical analyses

For all statistical data, values were derived from multiple measurements (from replicates of 3 or 5 experiments) and averaged, the standard deviations were evaluated using P-values from Student’s t-test (significance level α ≥ 0.05).

## 3. Results

### 3.1.1. *In silico* prediction of disordered regions of Eε34 TSP

The entire amino acid sequence of the wildtype as well as the Eε34 TSP were submitted to the online disordered region predictor and secondary structure prediction server known as PONDRs (Predictor of Natural Disordered Regions), a free online server (http://www.pondr.com/) and RaptorX (http://raptorx.uchicago.edu) **[31]** respectively. As shown in Figure 1, the Eε34 TSP showed higher disorder as compared to the wild-type protein. This is corroborated by the disordered regions as in indicated by red in the Raptor X prediction in Figure 2. In the Raptor X, predictions depicted amino acids that are in disordered regions as red and those in ordered region as blue.

**Figure 1.**
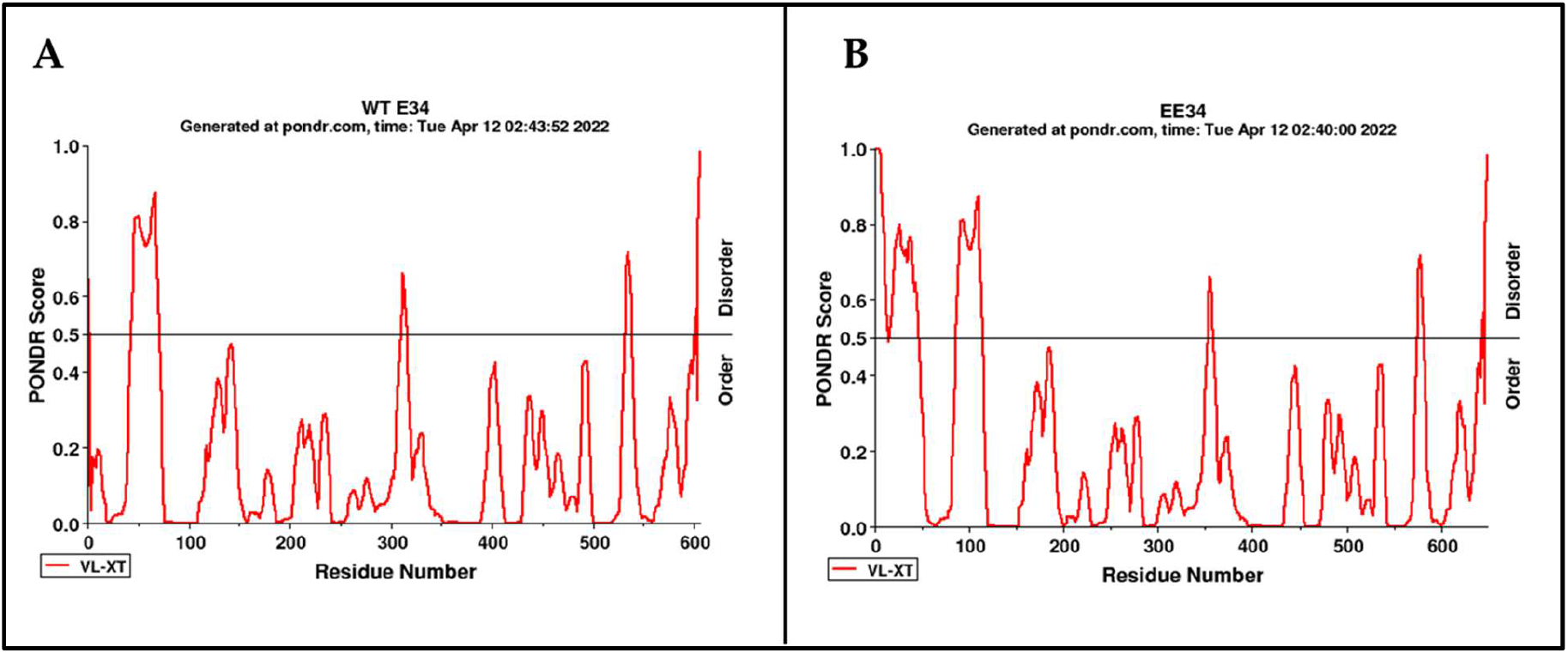
Disorder prediction of wild-type ε34 TSP (**A**) and extended ε34 TSP (**B**). Predictions were made using PONDRs (Predictor of Natural Disordered Regions), a free online server [http://www.pondr.com/]. To support this argument that the fusion peptide imparted disorder to the protein especially at the N-terminus, we analyzed for secondary structure disorder using the Raptor X. As shown in Figure 2. the amino acids that are predicted to have order (all sequences in blue color), and those that were disordered (all sequences in red color) are revealed.

**Figure 2.**
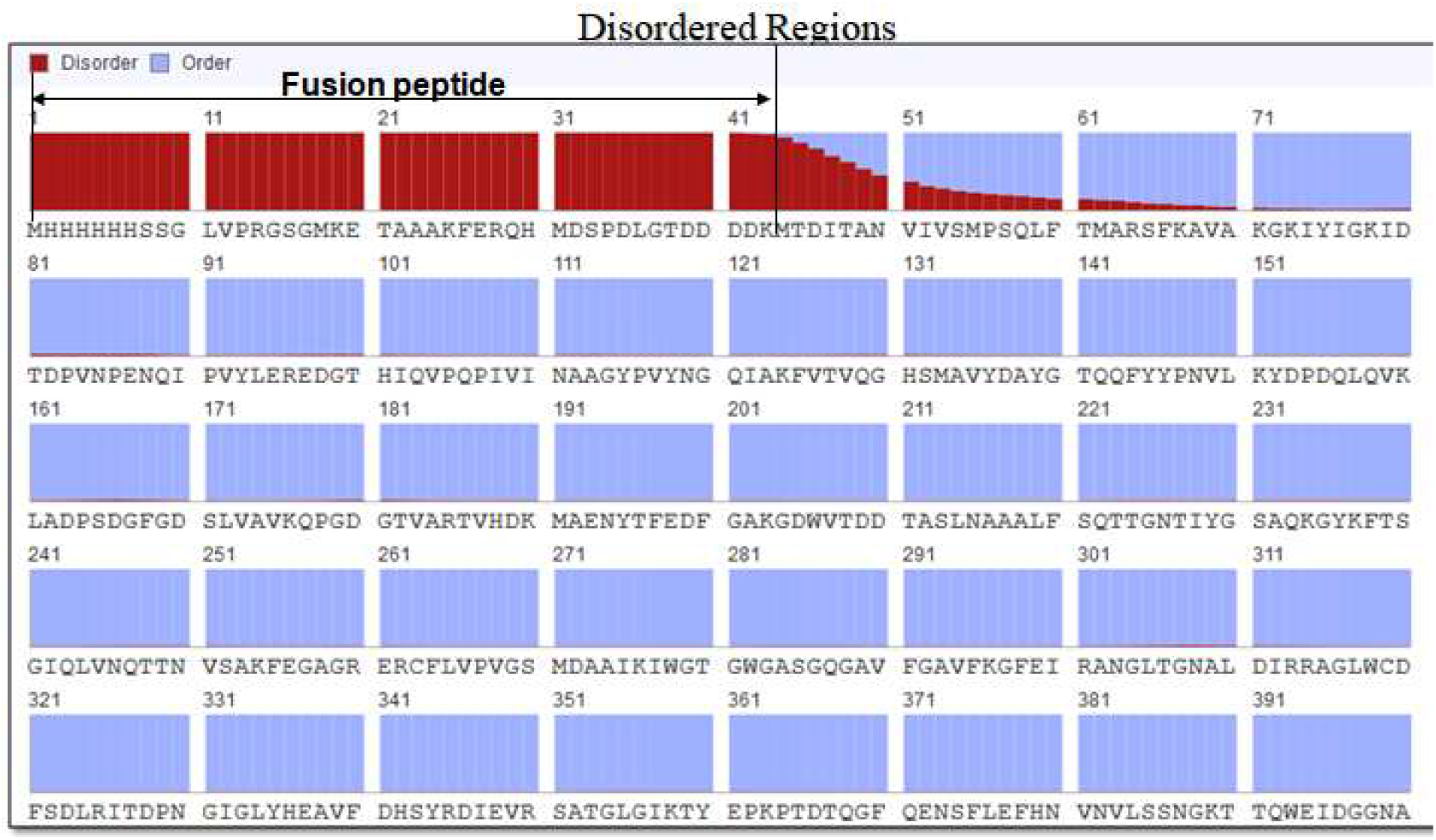
Secondary structure prediction by Raptor X shows the entire 43 amino acid sequence (the fusion peptide) to be completely disordered out of the first 400 amino acid sequence of the protein. With 100% disorder prediction, the 43 amino acid sequence seems to further disrupt the succeeding downstream sequences, the highest impact felt by the first 7 amino acids immediately following the fusion peptide (MTDITAN) which scored above 50% disorder. Disorder percentage tapered and faded at the 70th amino acid (A). The effect of fusion peptide on protein stability has been well documented **[32, 33]** but no answer exists on why most fusion peptide exert such influence on native protein’s stability. While we propose that the instability of our protein to proteinase K is partly due to the unstructured fusion tag, we do not doubt the fact that the very observed properties of the protein could be an intrinsic behavior of the protein under those conditions, hence further biophysical characterization is warranted

### 3.1.2. Proteinase K sites in wild-type ε34 TSP as compared to Eε34 TSP

As shown in Figure 3, the predicted PK sites on a denatured wild-type, as well as a native state ε34 TSP has been shown. Prediction were carried out using Raptor X server, where as the tertiary structure PK sites were predicted by peptidecutter and curated manually after identifying the secondary structure disordered regions.

**Figure 3.**
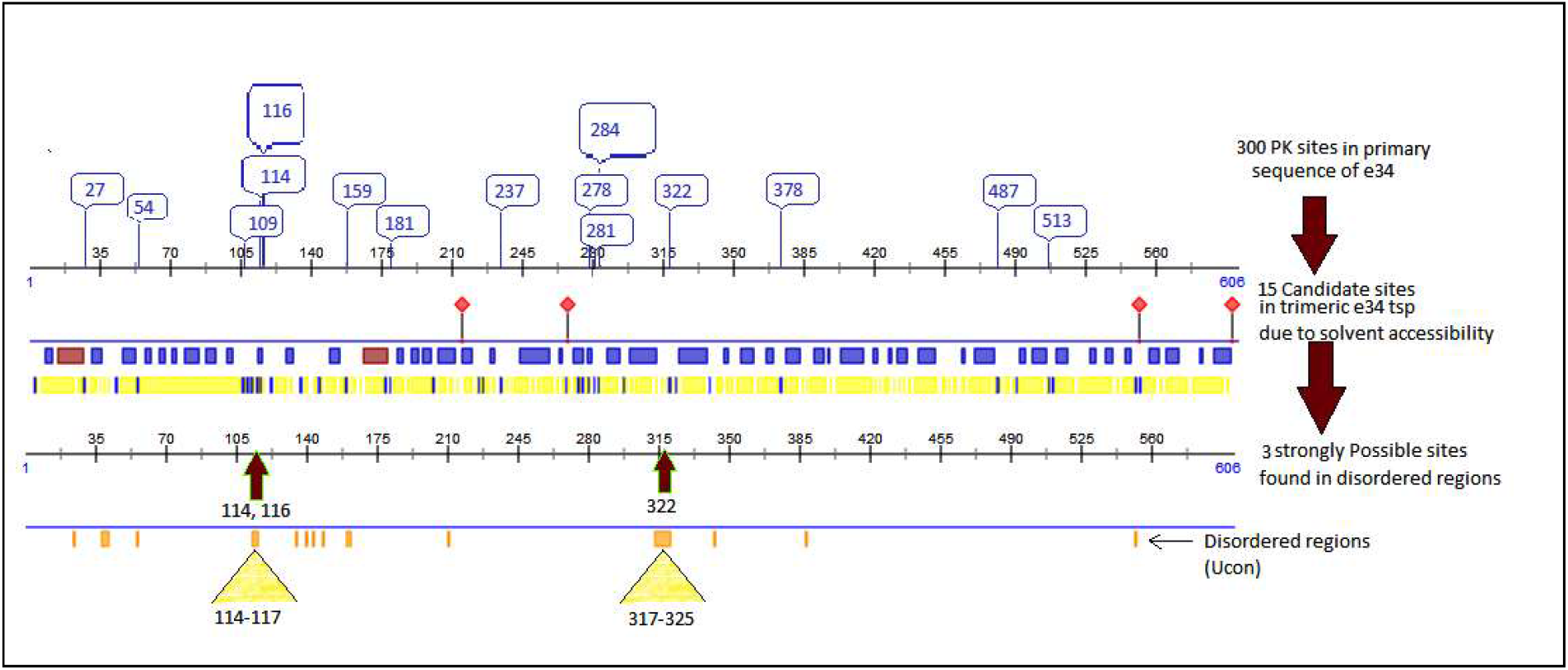
Diagrammatic illustration of PK site in the WT ε34 TSP. The red diamond depicts putative LPS binding sites of the TSP.

As illustrated the wildtype ε34 TSP contains 300 PK sites; however the Eε34 TSP contains 311 K sites (Figure not shown). In the quaternary conformation of the protein, the wild-type ε34 TSP is predicted to contain 3 sites as shown by the black arrows to be solvent accessible thus are highly likely to be accessible to PK, hence, presupposing that the protein is liable to be cleaved at these sites. In the case of Eε34 TSP, there are 14 predicted sites, and 11 of these sites are located on the 43 fusion peptide. The understanding of the possible cleavage sites especially on the native state of the protein is crucial guide to analyzing the peptide map generated after PK proteolysis of the protein.

### 3.2. Proteolytic analysis of Epsilon 34 Phage TSP indicates partial sensitivity to Proteinase K

#### 3.2.1. Partial Sensitivity of Eε34 TSP in 2% SDS to PK

Shown in Figure 4, however, the treatment of our Eε34 TSP in SDS to proteinase K at 37 °C for 6 h as depicted in lane 4 (EETUH) generated two protein fragments, one observed at 82 kDa and the other observed at 77 kDa, while the untreated trimers showed at the 161 kDa level. This observed significant migration difference between the untreated sample which migrated at 161 kDa and the treated samples which migrated at 82 kDa and 77 kDa possibly indicate the sensitivity of our protein to PK. When the treated samples were heat-unfolded to their monomeric state (Figure 4, lane 5, EETH), two monomeric bands were observed at 59 kDa and 25 kDa. In lane 6, the untreated and unheated matured samples migrated at the usual size, 123 kDa, but when treated to PK as in lane 8, a 77 kDa fragment is recorded, when the treated samples of the mature protein was subjected to heat unfolding, as in lane 9, we failed to observe any band, however, the untreated samples which were unfolded via heat recorded a monomeric band at 65 kDa as depicted in lane 7 of Figure 4. P22 TSP control, which was treated to PK showed complete resistance, registering a single band as depicted in lane 10, Figure 4. This led to our assumption that even though the trimeric Eε34 TSP shows sensitivity to PK, a much stable and protease resistant products resulting from the treatment were produced. This study demonstrates that Eε34 TSP is possibly liable to partial proteolytic digestion by proteinase K even at physiological temperatures of 37 °C in the presence of SDS (as depicted in Figure 4 lanes 4, 5 and 8). The present data points to cleavage sites that are possibly located interiorly of the trimeric protein, with possibility of cleaving a larger fragment off the TSP than we observed in the trypsin treatment.

**Figure 4.**
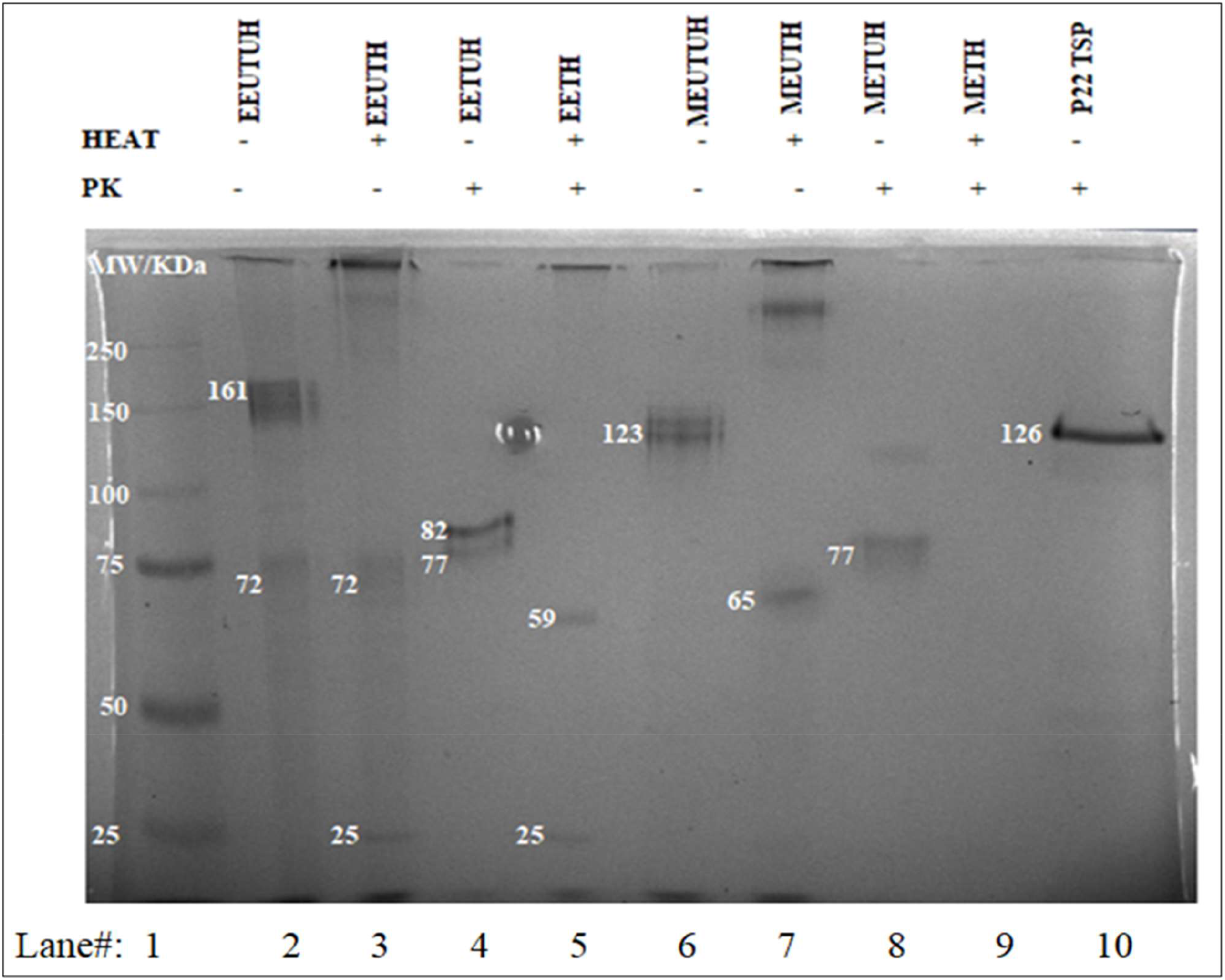
Native-PAGE analysis of Eε34 TSP in 2 % SDS treated to PK at physiological temperature. 0.5 mg/mL of Eε34 TSP and matured/rEK digested ε34 TSP (Mε34 TSP) in 2 % SDS were treated with proteinase K at 37 °C for 6 h. Samples were electrophoretically analyzed using 10 % polyacrylamide gel. Lane 1, Prestained Precision Protein Standard. Lane 2, Eε34 TSP untreated and unheated (EEUTUH). Lane 3, Eε34 TSP untreated but heated (EEUTH). Lane 4, Eε34 TSP treated but unheated (EETUH). Lane 5, Eε34 TSP treated and heated (EETH). Lane 6, matured ε34 TSP (rEK digested Eε34 TSP) treated but unheated (METUH). Lane 7, matured ε34 TSP (rEK digested Eε34 TSP) untreated but heated (MEUTH). Lane 8,matured ε34 TSP (rEK digested Eε34 TSP) treated but unheated (METUH). Lane 9, matured ε34 TSP (rEK digested Eε34 TSP) treated and heated (METH). Lane 10, P22 TSP treated but unheated.

#### 3.2.2. Eε34 TSP sensitivity test under increasing concentrations of enzyme

Eε34 TSP when unfolded to its monomeric state is 71.4 kDa, and seems to migrate at approximately 72 kDa on 7.5 % polyacrylamide gel. The sum of the molecular weight of the 43 amino acids fusion peptide and the last 65 amino acids molecular weight of the C-terminus (the trimerization domain) is approximately 11.9 kDa constituting the regions sensitive to PK, and if this molecular weight is subtracted from the 71.4 kDa molecular weight of the Eε34 TSP, a 59.5 kDa fragment is resulted. We propose that PK cleaves the N-terminally fused 43-amino acid fusion peptide and the last 65 amino acid fragment of the C-terminus, but unable to affect the head binding domain or the beta-helix domain of the protein (the region not sensitive to PK) as diagrammatically illustrated in **Figure 5**. The first 43 amino acids which is N-terminally fused to the protein is predicted to be completely unstructured (Figure 1 and 2). Hence, the unstructured nature of the fusion peptide makes it liable to degradation by PK.

**Figure 5.**
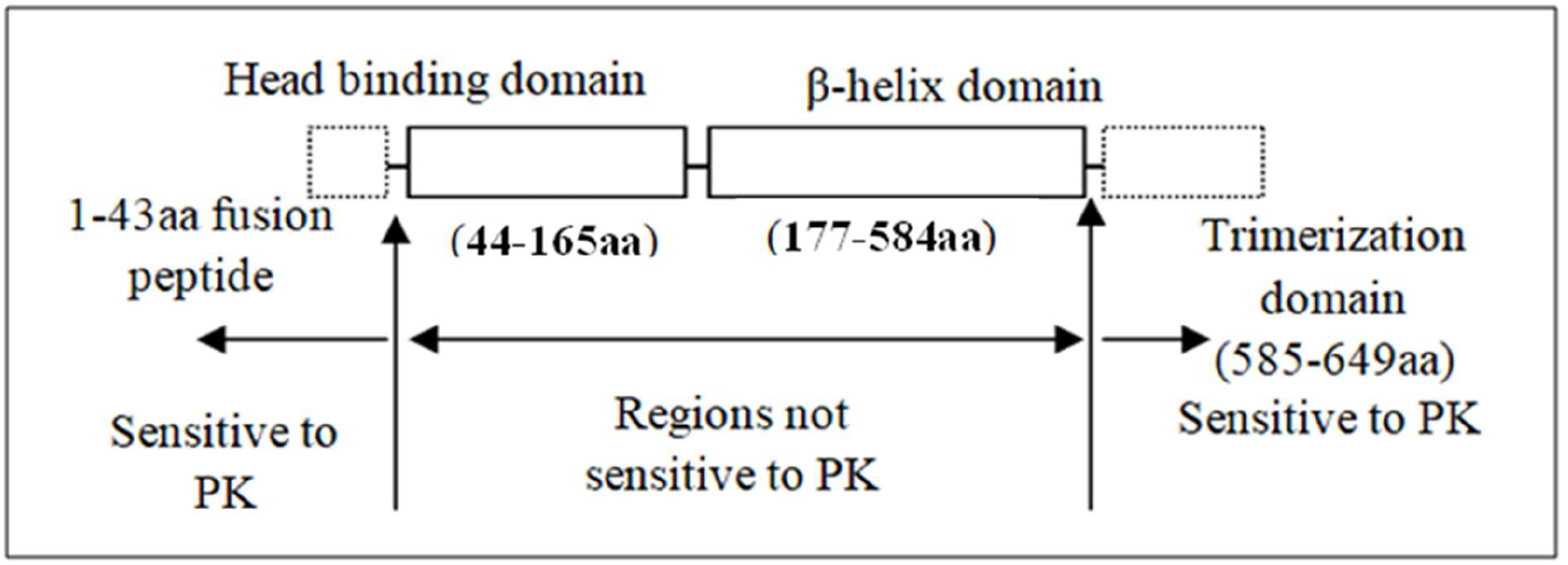
Diagrammatical depiction of the possible sensitive regions of trimeric Eε34 TSP to PK digestion (Created using Microsoft word).

Also, from solvent accessibility and disorder assessment data generated by the Raptor X, the entire fusion peptide scored high values and were completely disordered. In solvent accessibility heat maps generated by the Raptor X server, the protein registered high values for solvent accessibility at the positions S585 and I586, and these residues in the secondary structure assessesment is flanged at both sides by coils arising from sites A582, P583, and H584, I585, S587, V588 (Data not shown), however only sites A582 and I585 are PK cleavage sites, these all indicating a high possibility of PK accessing and binding to these sites to cleave the fragment off the protein.

The first 43 aa (amino acid) is the fusion peptide, and is predicted to be unstructured and labial, easily proteolyzed by PK (proteinase K). The sites A582 and I585 are the possible PK cleavable sites between the β-helix domain and the trimerization domain. Sum of the molecular weights of 43aa and 65 aa (i.e. 585-649 aa)=11.9 kDa. The difference in molecular weights between entire Eε34 TSP and PK sensitive domains = 59.5 kDa (i.e.(71.4 – 11.9) kDa) = Region resistant to PK at non-denaturing conditions.

#### 3.2.3. Testing the effect of increasing dose of PK to Eε34 TSP digestion

To understand if increasing the concentration of PK will affect the proteolysis of Eε34 TSP, varying concentrations of PK were administered to the protein (Figure 6). Digested samples were subjected to 7.5% native PAGE electrophoresis. As shown in Figure 6, increasing concentrations of PK did not affect the amount or size of the fragments generated.

**Figure 6.**
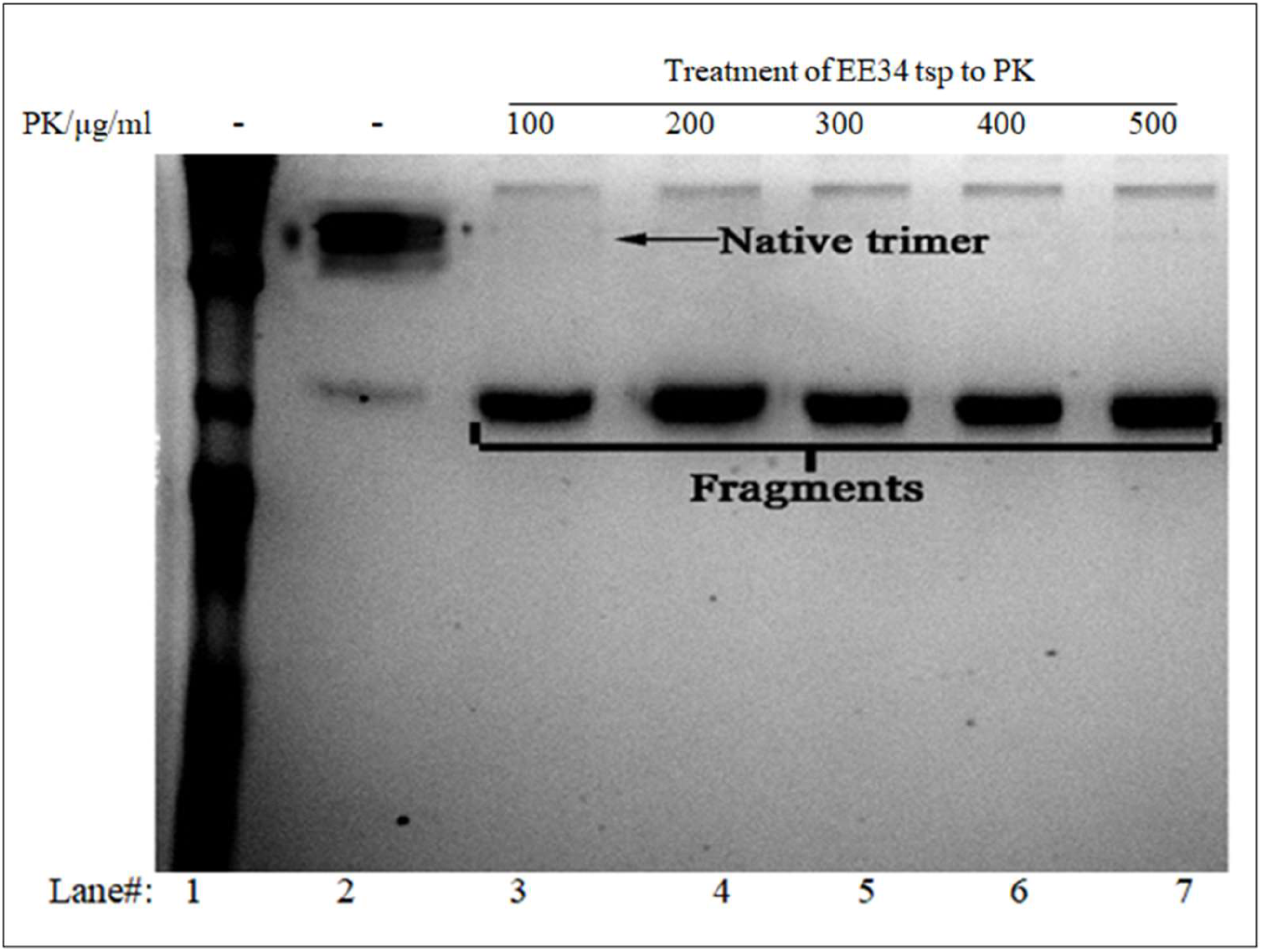
Prolonged enzymatic digestion of Eε34 TSP in varying concentrations of PK. 7.5 % Native PAGE gels analysis of resulting fragment arising from dose dependent treatment of the protein to PK. Lane 1, PPPS; Prestained Precision Protein Standard. Lane 2, Eε34 TSP unheated and untreated. Lane 3 to 7 are Eε34 TSP treated to varying concentration of PK.

#### 3.2.4. Incomplete inhibitory action of the protease inhibitor PMSF on Proteinase K

To gain insight into the ability of PMSF to inhibit the proteolytic activity of PK, we subjected sample of Eε34 TSP containing PMSF (10 µg/mL) to increasing dosages of PK. We observed that increasing concentrations of PK resulted in increased band intensities (Figure 7). While it was observed that all treatments (lanes3 to 7) produced fragments (FB) that migrated close to the monomeric size, it was clear nonetheless that trimeric species populated in high amounts in all samples as indicated by the trimeric band represented by the native band (NB) indicated in Figure 7. This trimeric band seemed to slightly increase with increasing concentrations of the enzyme (PK). This anomaly seems not to fit any feasible explanation given that approximately equal amount of TSP has been treated with the varying concentration of the enzyme. For instance, given that it was the incremental binding of PK to the TSP and yet lacking the proteolytic action due to the presence the protease inhibitor (PMSF) that led to the increasing trimeric band intensity, then the flaw will remain in the question, why did the binding of the enzyme not increase the migratory size of the trimers significantly? Another hypothesis could be that increasing concentrations of PMSF ensured that an increased amount of trimeric species were competitively protected from the protease, then if this argument is true, there should be decreasing intensities of the second fragment, which seems to hold true for lanes 3, 4, 5 and 7. The intensity of band in Lane 6 seems not to correlate with this deduction.

**Figure 7.**
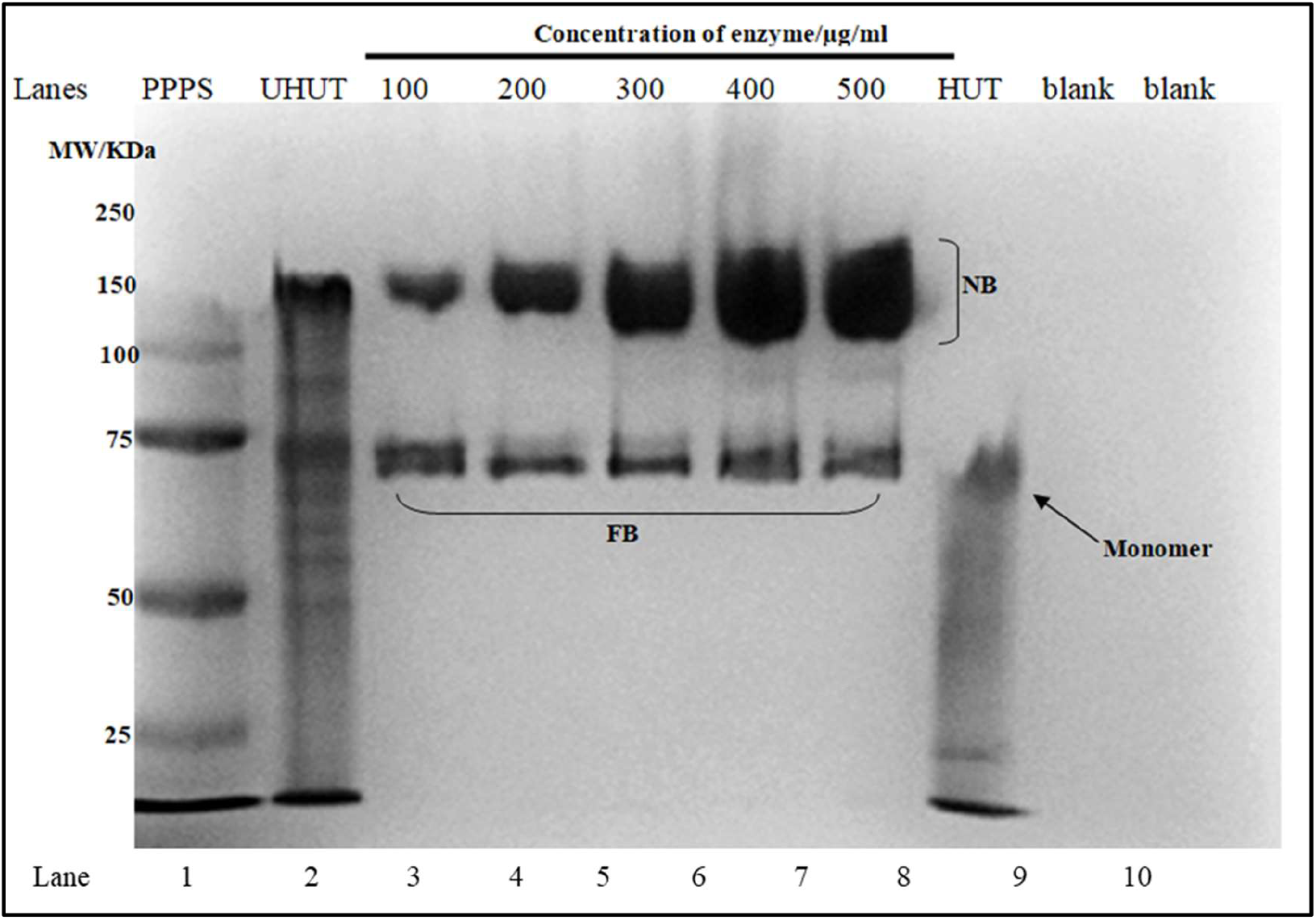
Proteolytic digestion of Eε34 TSP in 6 h using the enzyme proteinase K (PK), at incubation temperature of 37 °C and in the presence of PMSF.

Five samples of 1mg/ml of Eε34 TSP each was subjected to varying concentration of PK digest for 6 h under 37 °C and in the presence of PMSF. Afterward, samples were electrophoretically analyzed in a 7.5% SDS PAGE gel. In here, we loaded Lane 1, PPPS; Prestained Precision Protein Standard. Lane 2, Eε34 TSP; unheated and untreated (Eε34UHUT). Lane 3, Eε34 TSP; treated with 100µg of PK (Eε34UHT). Lane 4, Eε34 TSP; treated with 200 µg of PK(Eε34UHT). Lane 5, Eε34 TSP; treated with 300 µg of PK(Eε34UHT). Lane 6, Eε34 TSP; treated with 400 µg of PK(Eε34UHT). Lane, Eε34 TSP; treated with 500 µg of PK(Eε34UHT). Lane 8, Eε34 TSP; Heated and untreated (Eε34HUT). Lane 9, Blank, Lane 10, Blank.

#### 3.2.5. Quantitative analysis of incomplete inhibitory action of PMSF on Proteinase K

To better, understand the mechanism by which Eε34 TSP is digested by PK in the presence of PMSF as in Figure 7, the experiment was replicated and the densitometric values measured via imageJ. The mean values were used in plotting a chart as shown in Figure 8.

**Figure 8.**
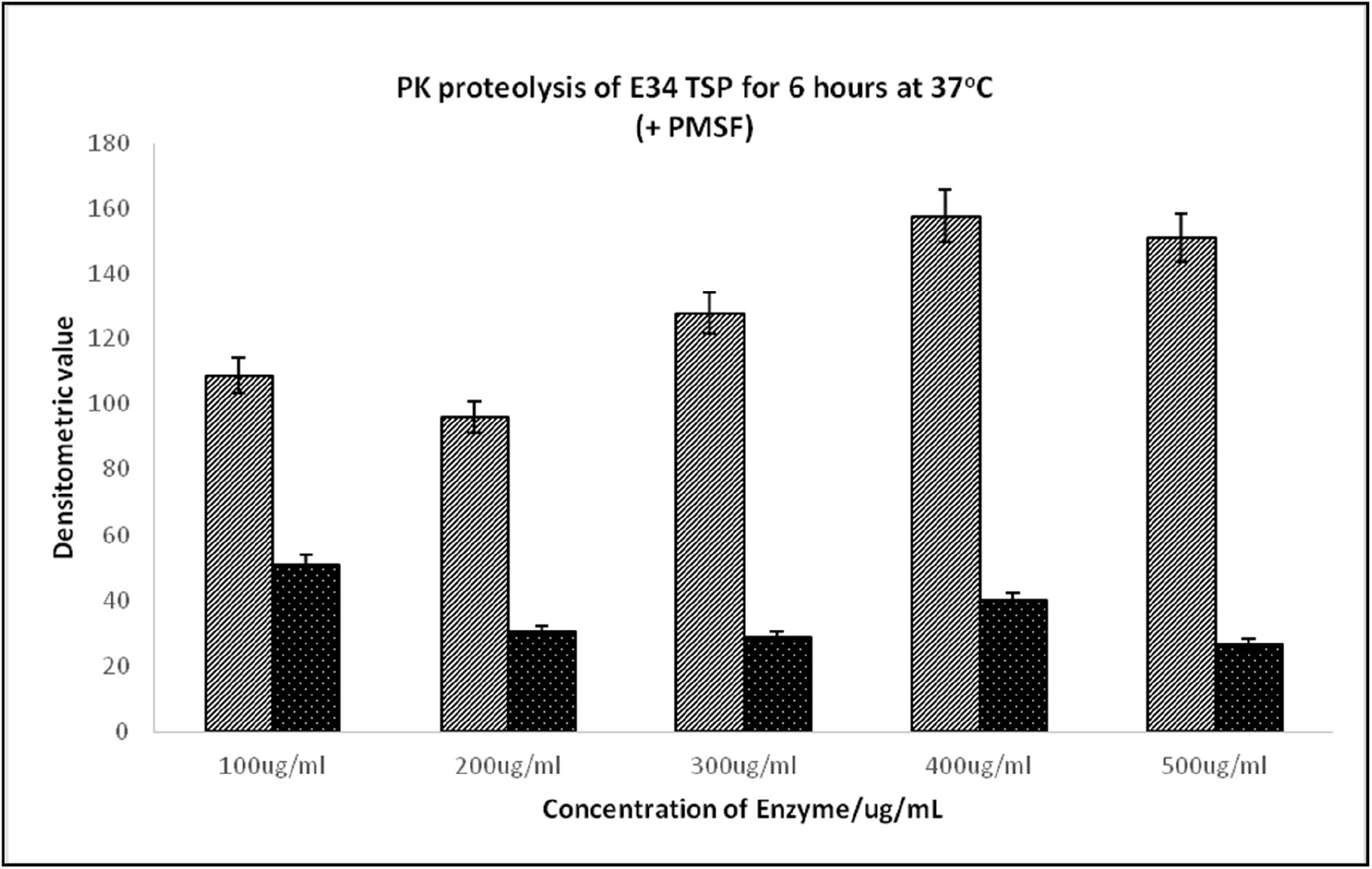
Limited proteolysis of Eε34 TSP in 6 h under the incubation temperature of 37 °C and in the presence of PMSF using PK Five samples of 1mg/ml of Eε34 TSP each were subjected to varying concentration of PK digest for 6 h under 37 °C and in the presence of PMSF. Afterward, samples were divided into two and electrophoretically analyzed in a 7.5% SDS PAGE gel. Replicates SDS PAGE gels were analyzed using densitometry. The values were averaged and plotted into a chart. Two bands designated as native band (NB, lighter columns) and fragment band (FB, the darker columns) were plotted.

The mean densitometric values for the trimeric species band (NB) were as following; 68.2, 96.2, 127.9, 157.9 and 151.2 representing the 100, 200, 300, 400 and 500µg/mL of PK treatment to samples respectively. While the fragment band (FB) registered the mean densitometric values 112.3, 91.8, 90.1, 101.3, and 87.8 for 100, 200, 300, 400 and 500 µg/mL PK treatment to Eε34 TSP samples respectively. While it is a possibility to deduce using this data, that, the increasing values as recorded in the NB were the increasing population of less stable trimeric species that were rescued in increasing amounts as PMSF increases, and yet the observed increase in trimeric band did not fully correlated with the FB band which represents the less stable species which were cleaved. The mean densitometric values for the fragment band (FB) for 100, 200, 300 and 500 µg/mL PK treatment seems to show a slight continued decrease in intensity, following our hypothesis that the less stable trimers were rescued/protected from proteolysis by increasing concentrations of the enzyme, however, the treatment 400µg/mL seems to behave as an outlier in this model.

### 3.3. Proteolytic digestion of Eε34 TSP at 70°C using PK

To investigate the effect of temperature on Eε34 TSP (Figure 9 and 10), we used PK digestion as a tool to scrutinize the unfolding kinetics of the protein. Here, we subjected Eε34 TSP to a high temperature of 70 °C (in a water bath preset at that temperature) for varying time points. Heating of samples were done without the addition of SDS, nor any other denaturants, in this way, it provided the platform to study the effect of heat alone at 70 °C on our TSP. Heated Eε34 TSP samples were cooled to room temperature, and then subjected to PK treatment. The electrophoretic mobilities of PK treated samples in all lanes 0, 2.5, 5, 10, 20, 30 and 60 (Figure 9) migrated to 77 kDa, a significant shift from the untreated Eε34 TSP migration of 161 kDa. No secondary fragments were observed in all treated samples.

**Figure 9.**
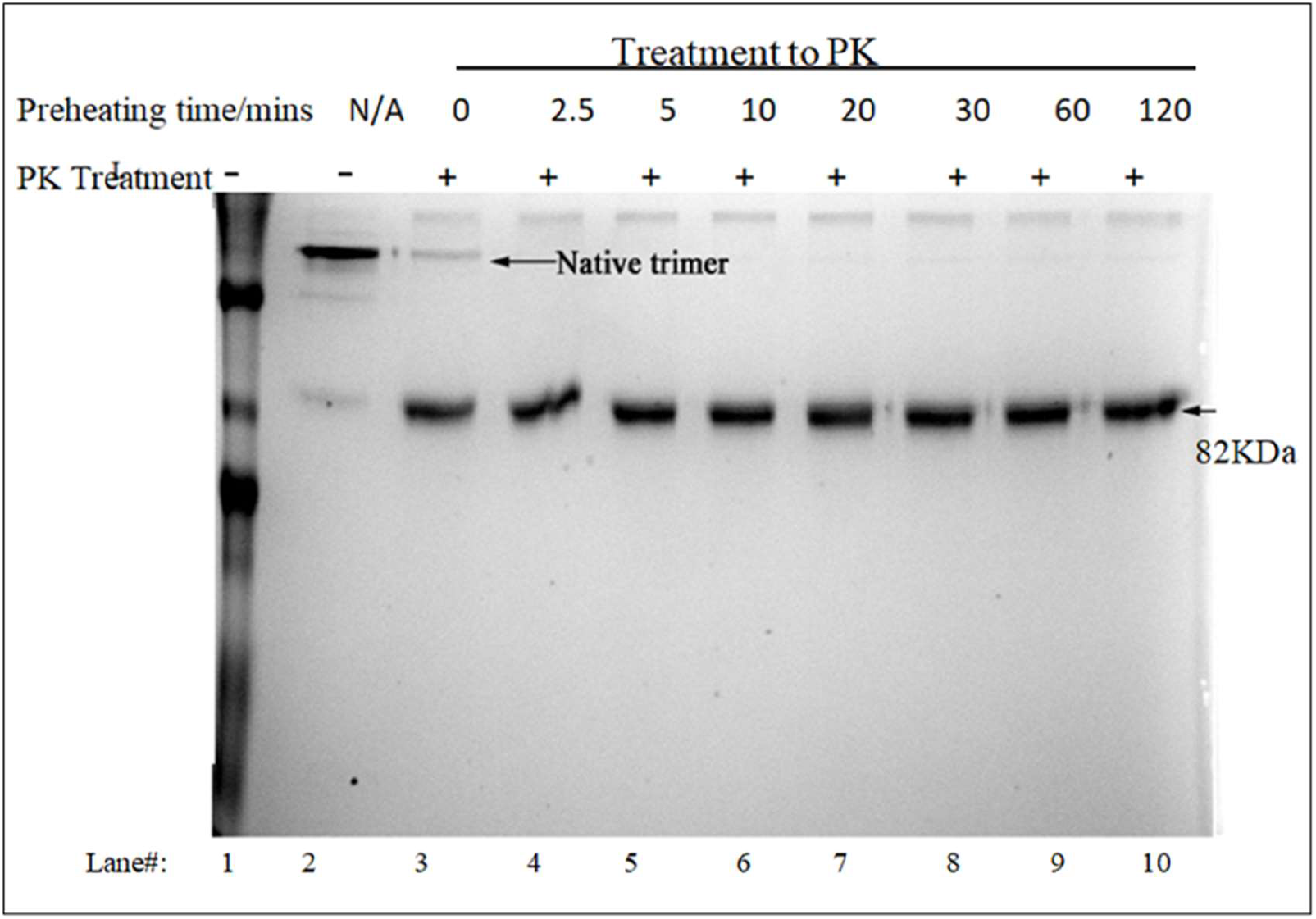
Native-PAGE analysis of PK proteolytic digestion of Eε34 TSP after 70 °C treatment. 0.5 mg/mL of Eε34 TSP samples were heated at 70 °C without the addition of SDS, then were withdrawn at set time points 0, 2.5, 5, 10, 15, 20, 30 and 60 min as indicated in lanes 3 to 9 respectively followed by PK digestion. Lane 2 served as unheated and untreated control. Lane 1 was PPPS (prestained perfect protein standard).

**Figure 10.**
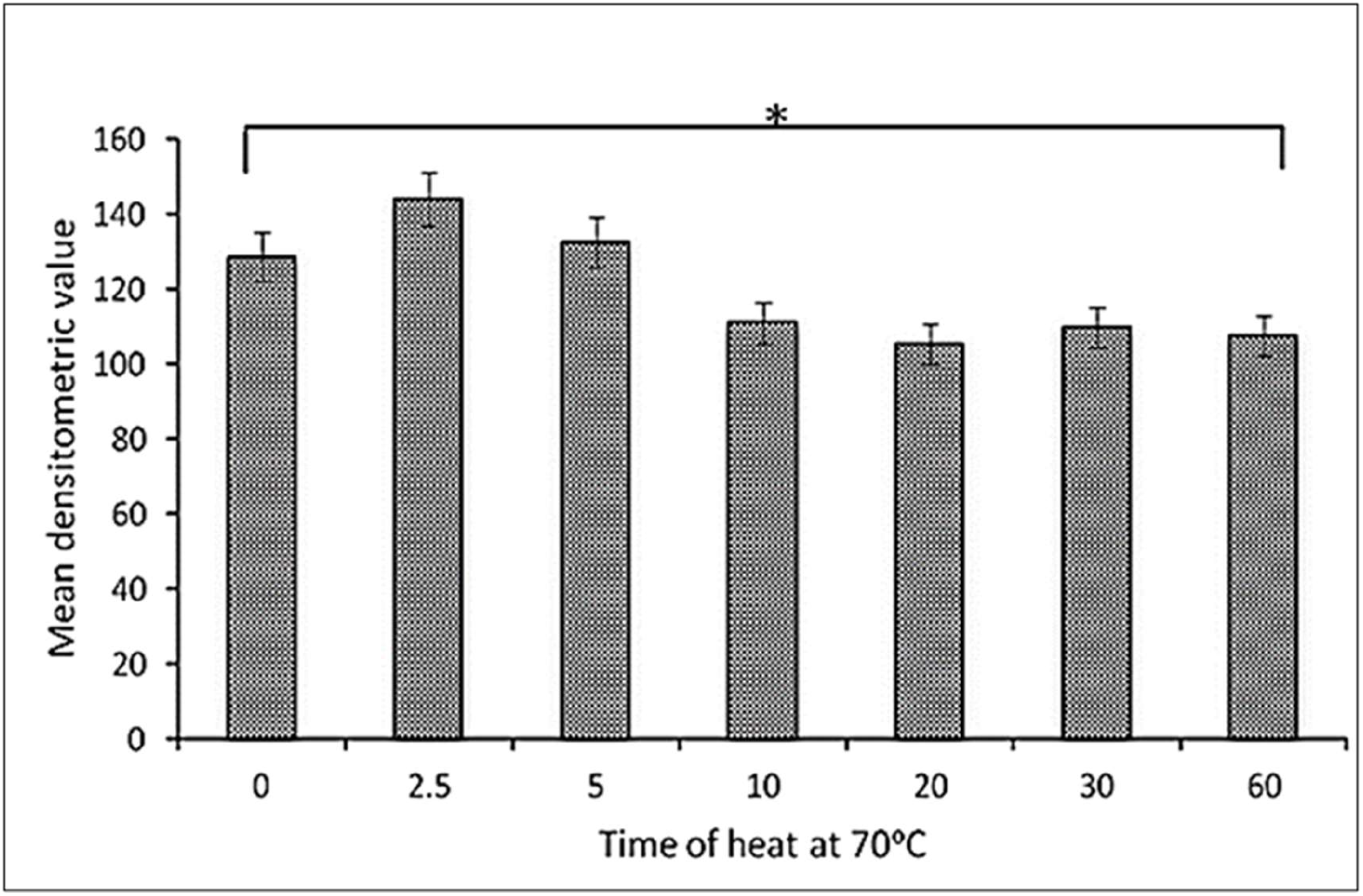
Bar plot of residual trimers after different time points of heating at 70 °C followed by PK digestion This chart compares residual Eε34 TSP fragments species resulting from the treatment of the protein to heat at 70 °C followed by PK digest. All data presented are derivative of triplicate experiments arising from Figure 9; all values represent the mean ±SEM (n = 3, p < 0.05).

As depicted in Figure 10, complementary to our observation (Figure 9), the plots of the mean densitometric values obtained illustrated almost similar values for some treatments, however, slight variability did exist. Beginning with a densitometric mean value of 128.58 at the 0-time point, the mean densitometric values that followed were as thus: 144.06, 132.58, 110.99, 105.52, 109.81 and 107.62 for time points 2.5, 5, 10, 20, 30 and 60 min respectively. Statistical differences were observed between means at p values less than 0.39 with the 2.5-minute treatment showing the highest difference. Within the limits of experimental errors, it is safe to argue for a continued marginal reduction density of bands with passing time, with the 60-minute treatment recording the lowest densitometric value of 107.62. This revelation might be a pointer to a marginal denaturation of some species of the protein at 70 °C with preponderance still existing in trimeric state but increasing incubation time meant continual accumulation of unfolded species which fell as substrate to PK complete digestion, hence the decreased intensities recorded at the 60^th^ minute.

### 3.4. Proteolytic digestion of Eε34 TSP at 80 °C using PK

#### 3.4.1. Qualitative analysis of Eε34 TSP treatment to PK after 80 °C incubation

In this experiment as depicted in Figure 11, it was noticed that a single fragment band was seen from lane 3 to lane 9, which represents, 0, 2.5, 5, 10, 20, 30 and 60 min of heating of samples at 80 °C before treatment with PK. The migration of the fragment band was recorded at 77 kDa, just slightly above the monomeric band which shows a migration of 7,1360 Da, as seen in lane 10 of Figure 11. The unheated and untreated samples treatment registered its band migration at the native trimeric size as indicated in lane 2. The fragment band migrated for all treated lanes at 77 kDa and is demonstrated to have a continued reduction in intensity as the time of heating at 80 °C increased. The faintest band was recorded at the 60^th^ minute mark as shown in lane 9 of Figure 11. Lane 10, which represents the heated but untreated samples registered a high-density band at the uppermost part of the gel, indicating aggregation, a slight band however can be noticed at the monomeric size, as indicated by the black arrow.

**Figure 11.**
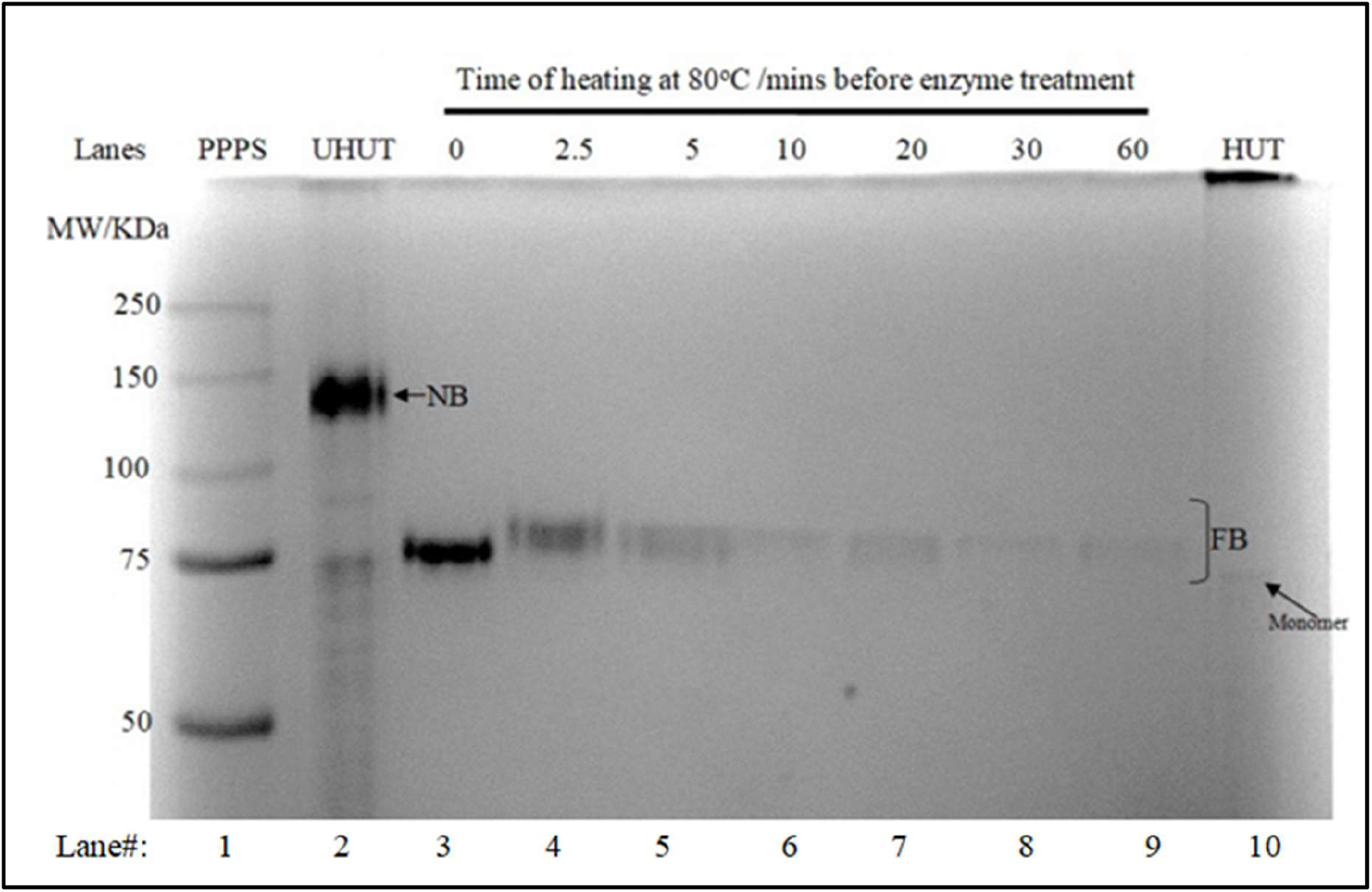
PK proteolysis of Eε34 TSP after 80 °C heat.

#### 3.4.2. Quantitative analysis of Eε34 TSP treatment to PK after 80 °C incubation

As demonstrated in Figure 12, we examined the unfolding rates as a function of time via the use of the proteolytic enzyme, PK. We quantitize and averaged the band intensities from three replica experiments. At zero time point, with a high mean densitometric value of 102.15, the proteolytic fragment band intensity decreased progressively with time to register a low of 0.14 densitometric value after 1 h of heating at 80 °C. The intervening time points 2.5, 5, 10, and 30 min registered densitometric values of 79.5, 62.3, 52.8, 32.9, and 16.4 respectively; showing that there was a progressive decrease in densities of the bands with time. Using a regressional analysis, a plot of these values seems to fit to an exponential curve with the equation; *[f*_*t*_*] = [f*_*0*_*]e*^*(−kt)*^; for which in this graph *[f*_*0*_*] =* 143.25 (theoretical, according to the curve), though practical value obtained was 102.15 for *[f*_*0*_] which represents the initial intensity (which equate to the concentration) of the proteolytic fragment band (FB), *k* = 0.105, *e =* mathematical constant = 2.718, Hence, to calculate for the resulting intensity or concentration of the proteolytic fragment remaining in solution at any time *t*; *[f*_*t*_*] = [*143.25*]e*^*(−0.105t)*^. For which *t =* time of heating (0, 2.5, 5, 10, 20, 30 and 60 min as indicated in this experiment) at 80 °C before treatment with PK. This equation shows the rate of decrease of the proteolytic fragment’s intensity with respect to time and can be conveniently used to estimate the concentration of the more heat resistant and studier truncated protein fragment at any given time. The principle is based on the fact that heating our protein unfolds it, however, the sequence of unfolding follows that the most labial and unstable regions unravels first, and the most stable regions melts last, hence giving the protease access to only the open regions of the protein at a sequential manner, hence those hitherto inaccessible sites are availed to the protease to bind to and cleave. In this work, a 30-min treatment of the samples to protease was sufficient to completely proteolyze a fully unfolded protein at the said concentration. *k* = the rate of proteolytic degradation of Eε34 TSP, which is also positively correlated to the rate of unfolding of the TSP, and was determined to be 0.105. In contrast to tryptic digest which registered k value of 0.072 **[3]**, PK treatment yielded a higher rate constant of 0.105, indicating a possibility of less stability of the truncated product formed via PK digestion as compared to the fragment produced in the tryptic digestion. This inference gains plausibility even further from published assessment of the importance of the trimerization domain of the P22 TSP to the stability of the protein. It has been demonstrated that the caudal fin of the P22 TSP, which translate here as possibly the last 65 amino acids sequence of the Eε34 TSP, the domain we propose to be digested by PK, functions as a molecular clamp in the P22 system **[34]**. This molecular clamp ensures thermostable subunit association in the native trimeric P22 TSP, therefore the possibility of similar role might not be farfetched in this Eε34 TSP. The truncated product has shown a high degree of thermal resistance, a known property of parallel b-helix domains [**35]**.

**Figure 12.**
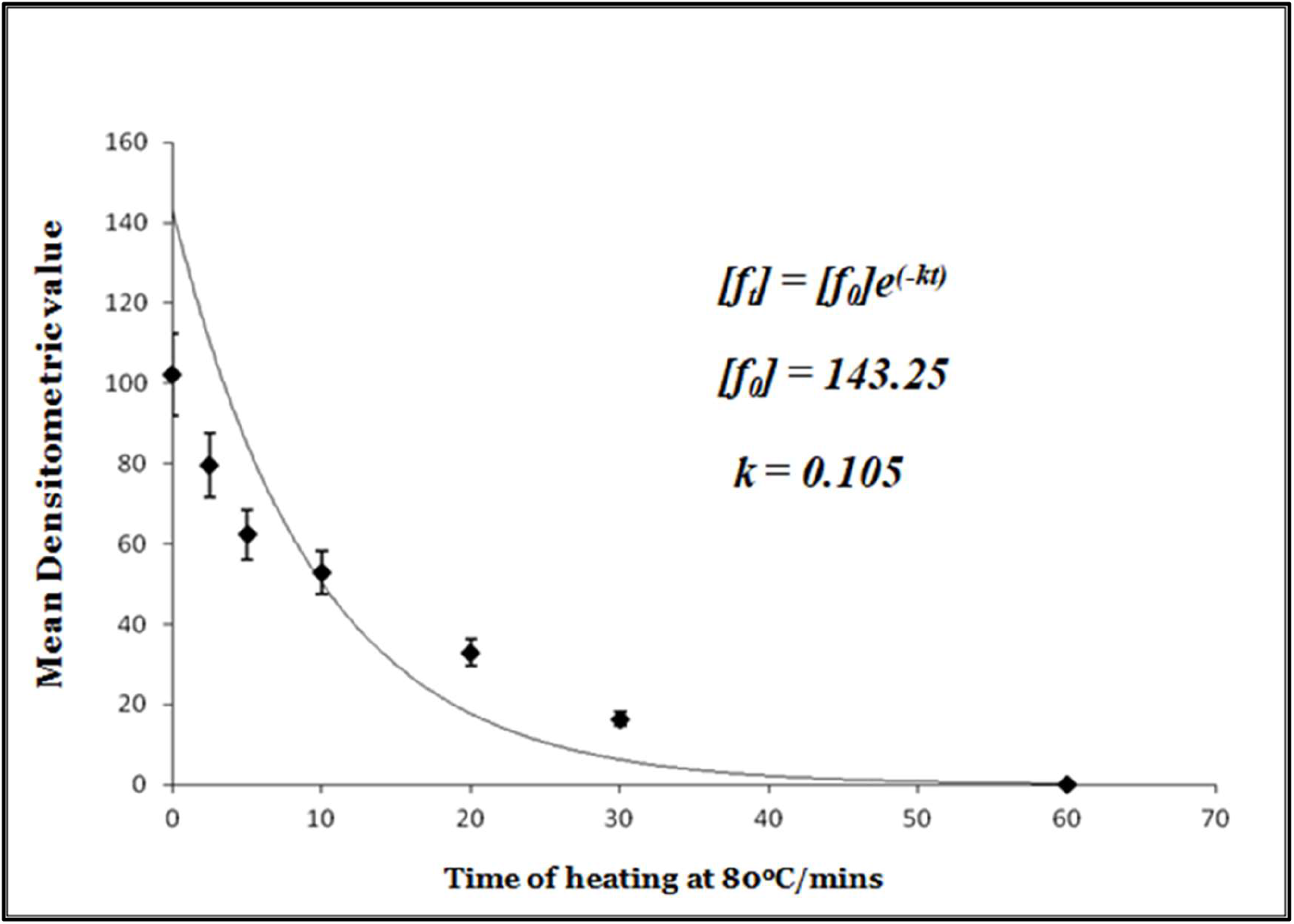
Quantitative assessment of unfolding kinetics of Eε34 TSP at 80 °C using PK. The mean densitometric value of three replicate experiments are presented; all values represent the mean ±SEM (n = 3, p < 0.05). These values were fitted into the unfolding curve, unfolding constant *(k*) was found to be 0.105. [*f*_*t*_]= concentration of Eε34 TSP at time *t*. [*f*_*0*_] = concentration of Eε34 TSP at the beginning of the experiment. *t*= time of heating.

### 4.0. Discussion

Trimeric protein in its 3D conformation presents complex topological surfaces and thus the native structural features of the Eε34 TSP have surfaces that are inaccessible to solvent thus rendering PK binding and subsequent proteolysis less effective. This sharply contrast the situation in a fully denature Eε34 TSP, because all the 311 proteinase K site along the length of the protein is readily available for digestion. For this reason, the understanding of the exact cleavage sites of these enzymes on Eε34 TSP primary sequence has provided means to speculate reliably the potential domains that are sensitive to the enzymes even at the native state topology. PK was predicted by PeptideCutty program to cleave Eε34 TSP in 311 sites, whereas it cleaves the wild-type ε34 TSP in 300 sites, and these sites are distributed unevenly throughout the entire sequence. However, upon secondary structure formation, subunit association, most of these sites become buried and shielded from the proteolytic enzyme, hence inferring proteolytic immunity to the protein.

An exhaustive review by Peter et al., 2003 [**36]** as well as other researchers in the field suggested a general sturdy nature of the β-helix domain in all TSPs and other proteins **[37, 38]**. As reported already when PK treated samples were heat-denatured, they migrated to approximately 59 kDa (Figure 4, lane 5), a fragment size that corresponded to the monomeric species of Eε34 TSP that is missing two components; the 43aa fusion peptide at the N-terminal region and the trimerization domain (caudal fin) that constitute the last 65 amino acid sequence of the protein. This hints at a possible cleavage at the linker region between the parallel β-helix and the triple β-prism of the Eε34 TSP. This is consistent with published findings concerning the robust nature of the β -barrel, which is known to be protease, heat and detergent resistant **[37, 38, 39, 40]**. The sum of the molecular weight of the 43 amino acid fusion peptide and the last 65 amino acids molecular weight of the C-terminus is 11880 Da, and if this molecular weight is subtracted from the 71390 Da molecular weight of the Eε34 TSP, a 59510 Da fragment is resulted. Although, this method is too crude to propose a conclusion, yet it is suggestive of a possibility that PK might have cleaved the N-terminally fused 43-amino acids fusion peptide and the last 65 amino acids fragment of the C-terminus (trimerization domain), but unable to affect the head binding domain or the beta-helix domain of the protein. As a control, P22 TSP and matured ε34 TSPs were also treated with PK. While it was obvious from the band migration that Mε34 TSP was sensitive to PK (Figure 4, lane 6), P22 TSP indicated no sensitivity at all to this enzyme (Figure 4, lane 8). The treatment of Mε34 TSP resulted in a protein fragment migrating at approximately 77 kDa while P22 TSP still maintained its usual trimeric electrophoretic mobility at the size of 126 kDa on 7.5 % polyacrylamide gel which corresponds to a 216 kDa in its actual molecular weight.

## Conclusion

Given the characteristics of Eε34 TSP observed, in this study, it indicates that ε34 TSP although previously shown to be resistant to trypsin, it is susceptible to proteinase K, at non-denaturing conditions. Further analysis of the fragments indicated that a studier and more heat resistant fragment was generated after proteinase K treatment, and this fragment seems to be resistant to heat even at elevated temperatures up to 70 °C. Nonetheless, the resulting fragment denatures at 80 °C or higher. The migration of the monomeric species of the truncated product from PK treatment showed a characteristic match for Eε34 TSP missing the fusion peptide and the trimerization domain. The topology of the predicted trimeric protein shows a flexible region at two PK sites located right between the LPS-binding central domain and the trimerization domain, and this could be a prove that PK binds to and degrade this region of the protein as well as the fusion peptide to produce a truncated product devoid ofthese two regions. Interestingly, further analysis of this truncated product shows a domain emerged to be resistant to SDS and to heat at 70 °C and hence must necessarily be a compact and sturdy product.

This study, apart from providing the structural sculpture of the Eε34 TSP, we showed that the unfolding kinetics of this protein can be probe via protease, heat, and detergent combinations. These simple tools can provide valuable information about unknown protein as regarding structural flexibility or compactness. Here, Eε34 TSP could be regarded as a typical example of hybrid molecule of which an intrinsically disordered protein (IDPs), that is the fusion peptide and structurally compact or ordered domains (that is the matured protein devoid of the fusion peptide) are married. The characteristic features observed in this study of Eε34 TSP for which the 43 amino acids fusion peptide is labial imparts this property of IDPs to a rather compact and highly organized architecture formed from very structured domains of well-defined three-dimensional folds arising from the wild-type protein. IDPs engage in most cellular functions **[41, 42]** in which structural plasticity presents functional advantage in terms of binding elasticity. Thus this study has presented a novel method to produce IDPs modules for classroom research, for bio-engineering purposes, or to understand certain peculiar nature of some proteins that are blamed for the etiology of some disease caused by IDPs.

## Statistical analyses

For all statistical data, values were derived from multiple measurements (from replicate of 3 or 5 experiments) and averaged, the standard deviations were evaluated using P values of Student’s *t* test (two tailed, two samples of unequal variance, significance level α ≥ 0.50).

## Acknowledgements

We thank the department of Microbiology, Faculty of Science, Mathematics and Technology, Alabama State University for timely support with supplies and quick responses to repairs of laboratory equipment.

## Ethical Approval

This article does not contain any studies with human participants or animals performed by any of the authors.

## List of Abbreviations Abbreviations

µL: micro liter
µM: micro molar
AGG: Aggregates
Da: Dalton
Native-PAGE: Native Polyacrylamide gel electrophoresis Eε34 TSP Extended Epsilon 34 tailspike protein
Ek: Enterokinase
ε34: TSP Epsilon 34 tailspike protein
Mε34: TSP Matured (enterokinase digested) ε34 TSP
HCl: Hydrochloric Acid
IDPs: Intrinsically Disordered Proteins
KDa: Kilo Daltons
mL: milliliter
LPS: Lipopolysaccharides
M: Molarity
Min: Minutes
mM: milli-molar
MM: Monomers
Mr, MW: Molecular Weight
N/A: Not applicable
NB: Native band
OD: Optical Density
PBS: Phosphate Saline Buffer
PK: Proteinase K
PMSF: PhenylMethanesulfonyl fluoride
PONDR: Predictor of Natural Disordered Regions
PPPS: Pre-stained perfect Protein Standard
rpm: Revolution per minute
SDS PAGE: Sodium Dodecyl Sulfate Polyacrylamide Gel Electrophoresis
SDS: Sodium Dodecyl Sulfate
SEM: Standard Error of Mean
TH: Treated and Heated samples of protein
TSP: Tailspike protein
TUH: Treated but unheated
UTH: Untreated but heated
UTUH: Untreated and Unheated

## References

1 Gildea, L., Ayariga, J.A. and Villafane, R., 2021. P22 Phage Shows Promising Antibacterial Activity Under Pathophysiological Conditions.

2 Centers for Disease Control and Prevention (CDC). National Salmonella Surveillance Overview. Atlanta, Georgia: US Department of Health and Human Services, CDC, 2011. http://www.cdc.gov/nationalsurveillance/PDFs/NationalSalmSurveillOverview_508.pdf (accessed 10/25/2012

3 Ayariga JA, Gildea L, Villafane R. ε34 Phage Tailspike Protein is Resistant to Trypsin and Inhibits Salmonella Biofilm Formation. Enliven:Microb Microbial Tech. 2022; 9(2): 002.

4 Ayariga, J.A., Abugri, D., Amrutha, B. and Villafane, R., 2022. Capsaicin potently blocks Salmonella typhimurium invasion of Vero cells. bioRxiv.

5 Gildea, L.; Ayariga, J.; Ajayi, O.; Xu, J.; Villafane, R.; Samuel-Foo, M. Cannabis sativa CBD Extract Shows Promising Antibacterial Activity against S. typhimurium and S. newington. Preprints 2022, 2022030367 (doi: 10.20944/preprints202203.0367.v1).

6 Gildea, L.; Ayariga, J.; Abugri, J.; Villafane, R. Phage Therapy: A Potential Novel Therapeutic Treatment of Methicillin-Resistant Staphylococcus aureus. Preprints 2021, 2021110397 (doi: 10.20944/preprints202111.0397.v1)

7 Maura, D. and Debarbieux, L., 2011. Bacteriophages as twenty-first century antibacterial tools for food and medicine. Applied microbiology and biotechnology, 90(3), pp.851–859.

8 Kropinski, A.M., Sulakvelidze, A., Konczy, P. and Poppe, C., 2007. Salmonella phages and prophages—genomics and practical aspects. Salmonella, pp.133–175.

9 Ayariga J. A., Gildea L, Wu H, Villafane R (2022) The ε34 Phage TSP: An In vitro Characterization, Structure Prediction, Potential Interaction with S. newington LPS and Cytotoxicity Assessment to Animal Cell Line. J Clin Trials. S14:002

10 Joseph A. Ayariga, Karthikeya Venkatesan, Robert Ward, Hongzuan Wu, Doba Jackson, and Robert Villafane (2018) Initiation of P22 Infection at the Phage Centennial, Frontiers in Science, Technology, Engineering and Mathematics, Volume 2, Issue 2, 64–81

11 Harada, Kenji, Mitsuo Kameda, Mitsue Suzuki, and Susumu Mitsuhashi. “Mutant of Salmonella phage epsilon 34 with loss of converting ability.” Japanese journal of microbiology 8, no. 4 (1964): 125–130.

12 Zayas, M.V. and Villafane, R., 2007. The tailspike protein of the Salmonella phage epsilon 34.

13 Harada, K., Kameda, M., Suzuki, M. and Mitsuhashi, S., 1964. DRUG RESISTANCE OF ENTERIC BACTERIA III. FR (TC) Acquisition of Transferability of Nontransmissible R (TC) Factor in Cooperation With F Factor and Formation of. Journal of bacteriology, 88(5), pp.1257–1265.

14 Williams, J., Venkatesan, K., Ayariga, J.A., Jackson, D., Wu, H. and Villafane, R., 2018. A genetic analysis of an important hydrophobic interaction at the P22 N-terminal domain. Archives of virology, 163(6), pp.1623–1633.

15 Ayariga, J.A. and Villafane, R., 2021. Single Amino Acid Change Mutation in the Hydrophobic Core of the N-terminal Domain of P22 TSP affects the Proteins Stability. bioRxiv.

16 Roach, D.R. and Donovan, D.M., 2015. Antimicrobial bacteriophage-derived proteins and therapeutic applications. Bacteriophage, 5(3), p.e1062590.

17 Ayariga JA, Gildea L, Villafane R. Extended ε34 Phage TSP Renatures after Urea-Acid Unfolding. Enliven: Microb Microbial Tech. 2022; 9(1): 001.

18 Reich, L., Becker, M., Seckler, R. and Weikl, T.R., 2009. In vivo folding efficiencies for mutants of the P22 tailspike β-helix protein correlate with predicted stability changes. Biophysical chemistry, 141(2-3), pp.186–192.

19 Yoder, M.D. and Jurnak, F., 1995. The parallel β helix and other coiled folds. The FASEB journal, 9(5), pp.335–342.

20 Gazit, E., 2005. Mechanisms of amyloid fibril self-assembly and inhibition: Model short peptides as a key research tool. The FEBS journal, 272(23), pp.5971–5978.

21 Rochet, J.C. and Lansbury Jr, P.T., 2000. Amyloid fibrillogenesis: themes and variations. Current opinion in structural biology, 10(1), pp.60–68.

22 Makin, O.S. and Serpell, L.C., 2005. Structures for amyloid fibrils. The FEBS journal, 272(23), pp.5950–5961.

23 Baxa, U., Steinbacher, S., Miller, S., Weintraub, A., Huber, R. and Seckler, R., 1996. Interactions of phage P22 tails with their cellular receptor, Salmonella O-antigen polysaccharide. Biophysical journal, 71(4), pp.2040–2048.

24 Yoder, M.D., Lietzke, S.E. and Jurnak, F., 1993. Unusual structural features in the parallel β-helix in pectate lyases. Structure, 1(4), pp.241–251.

25 Mótyán JA, Tóth F, Tőzsér J. Research applications of proteolytic enzymes in molecular biology. Biomolecules. 2013 Nov 8;3(4):923–42. doi: 10.3390/biom3040923. PMID: 24970197; PMCID: PMC4030975.

26 Barrett, A.J. and McDONALD, J.K., 1986. Nomenclature: protease, proteinase and peptidase. Biochemical Journal, 237(3), p.935.

27 Li, Q., Yi, L., Marek, P. and Iverson, B.L., 2013. Commercial proteases: present and future. FEBS letters, 587(8), pp.1155–1163.

28 Gill, J.J., Wang, B., Sestak, E., Young, R. and Chu, K.H., 2018. Characterization of a novel Tectivirus phage toil and its potential as an agent for biolipid extraction. Scientific reports, 8(1), pp.1–11.

29 Bosch, B.J., Van der Zee, R., De Haan, C.A. and Rottier, P.J., 2003. The coronavirus spike protein is a class I virus fusion protein: structural and functional characterization of the fusion core complex. Journal of virology, 77(16), pp.8801–8811.

30 Ebeling W, Hennrich N, Klockow M, Metz H, Orth HD, Lang H (August 1974). “Proteinase K from Tritirachium album Limber”. Eur. J. Biochem. 47 (1): 91–7. doi:10.1111/j.1432-1033.1974.tb03671.x. PMID 4373242.

31 Källberg, M., Margaryan, G., Wang, S., Ma, J. and Xu, J., 2014. RaptorX server: a resource for template-based protein structure modeling. In Protein structure prediction (pp. 17–27). Humana Press, New York, NY.

32 Reed, M.L., Yen, H.L., DuBois, R.M., Bridges, O.A., Salomon, R., Webster, R.G. and Russell, C.J., 2009. Amino acid residues in the fusion peptide pocket regulate the pH of activation of the H5N1 influenza virus hemagglutinin protein. Journal of virology, 83(8), pp.3568–3580.

33 Singh, P., Sharma, L., Kulothungan, S.R., Adkar, B.V., Prajapati, R.S., Ali, P.S.S., Krishnan, B. and Varadarajan, R., 2013. Effect of signal peptide on stability and folding of Escherichia coli thioredoxin. PloS one, 8(5), p.e63442.

34 Kreisberg JF, Betts SD, Haase-Pettingell C, King J. 2002. The interdigitated beta-helix domain of the P22 TSP acts as a molecular clamp in trimer stabilization. Protein Sci 4:820

35 Bradley P, Cowen L, Menke M, King J, Berger B. 2001. BETAWRAP: Successful prediction of parallel beta-helices from primary sequence reveals an association with many microbial pathogens. Proc Natl Acad Sci USA 98:148119–148224

36 Peter R. Weigele, Eben Scanlon, Jonathan King. 2003. Homotrimeric, beta-stranded viral adhesions and Tail proteins. Journal of Bacteriology. 185.14. 4022–4030

37 Junker M, Schuster CC, McDonnell AV, Sorg KA, Finn MC, Berger B, Clark PL. 2006. Pertactin beta-helix folding mechanism suggests common themes for the secretion and folding of autotransporter proteins. Proceedings of the National Academy of Sciences of the United States of America 103:4918–4923

38 Sauer RT, Krovatin W, Poteete AR, Berget PB. 1982. Phage P22 tail protein: Gene and amino acid sequence. Biochemistry 21:5811–5815

39 Zayas M, Villafane R. 2007. Identification of the Salmonella phage ε34 tailspike gene. Gene 386:211–217

40 Steinbacher S, Seckler R, Miller S, Steipe B, Huber R, Reinemer P. 1994. Crystal structure of P22 TSP: interdigitated subunits in a thermostable trimer. Science 265:383–386.

41 Christopher JO, Vladimir NU, Keith AD, Lukasz K. 2019. Introduction to intrinsically disordered proteins and regions. Academic Press 1–34

42 Jane DH. 2016. Making Sense of Intrinsically Disordered Proteins. Biophysical Journal 110:1013–1016

